# Keratin filament mechanics and energy dissipation are determined by metal-like plasticity

**DOI:** 10.1101/2022.11.05.515302

**Authors:** Charlotta Lorenz, Johanna Forsting, Robert W. Style, Stefan Klumpp, Sarah Köster

## Abstract

Cell mechanics is determined by an intracellular biopolymer network, including intermediate filaments that are expressed in a cell-type specific manner. A prominent pair of intermediate filaments are keratin and vimentin as the epithelial-to-mesenchymal transition is associated with a switch from keratin to vimentin. The transition coincides with a change in cellular mechanics, and thus dynamic properties of the cells. This observation raises the question of how the mechanical properties already differ on the single filament level. Here we use optical tweezers and a computational model to compare the stretching and dissipation behavior of the two filament types. We find that keratin and vimentin filaments behave in opposite ways: keratin filaments elongate, but retain their stiffness, whereas vimentin filaments soften, but retain their length. This finding is explained by fundamentally different ways to dissipate energy: viscous subunits sliding within keratin filaments and non-equilibrium *α* helix unfolding in vimentin filaments.

## Introduction

Biological cells possess an astounding composite materials system, the so-called cytoskeleton, which ensures mechanical integrity and stability and is responsible for active processes such as cell division and migration. Three families of biopolymers – actin filaments, microtubules and intermediate filaments – together with passive cross-linkers and active molecular motors form interpenetrating networks within cells,^1,2^ which adapt precisely to the mechanical needs and functions of each cell type. In contrast to actin and tubulin, intermediate filament proteins are expressed in a cell-type specific manner,^3–5^ making them ideal candidates for cells to adapt their mechanical properties.^6^ A prominent example of differential expression of intermediate filament proteins is the epithelial-to-mesenchymal-transition, ^7–13^ which occurs during cancer metastasis, embryogenesis^14^ and wound healing.^15^ These processes have in common that stationary, strongly interconnected epithelial cells change their phenotype to highly motile mesenchymal cells. Interestingly, epithelial cells typically express the intermediate filament protein keratin, whereas mesenchymal cells express vimentin.

We have recently shown that already on the single filament level keratin 8/18 filaments are softer and exhibit a very different force-strain behavior than vimentin filaments:^16^ Keratin filaments exhibit a nearly linear increase in force up to a strain of 0.7 after which filaments stiffen. Vimentin filaments are stiffer for strains for up to 0.15 and exhibit a plateau-like regime for strains between 0.15 and 0.8, in which the force barely increases. For strains larger then 0.8, vimentin filaments stiffen as well. These mechanical properties of intermediate filament proteins are closely related to their molecular architecture: ^17–19^ The monomers consist of three *α*-helical domains flanked by intrinsically disordered head and tail domains as sketched in Fig. 1a. ^20^ In case of keratin and vimentin filaments, 16 or 32 monomers, respectively, associate laterally in a stepwise manner to form unit-length filaments (ULFs).^3,20,21^ ULFs associate longitudinally to form filaments, thus resulting in an array of monomers arranged laterally and longitudinally and connected by electrostatic and hydrophobic interactions.

**Figure 1:**
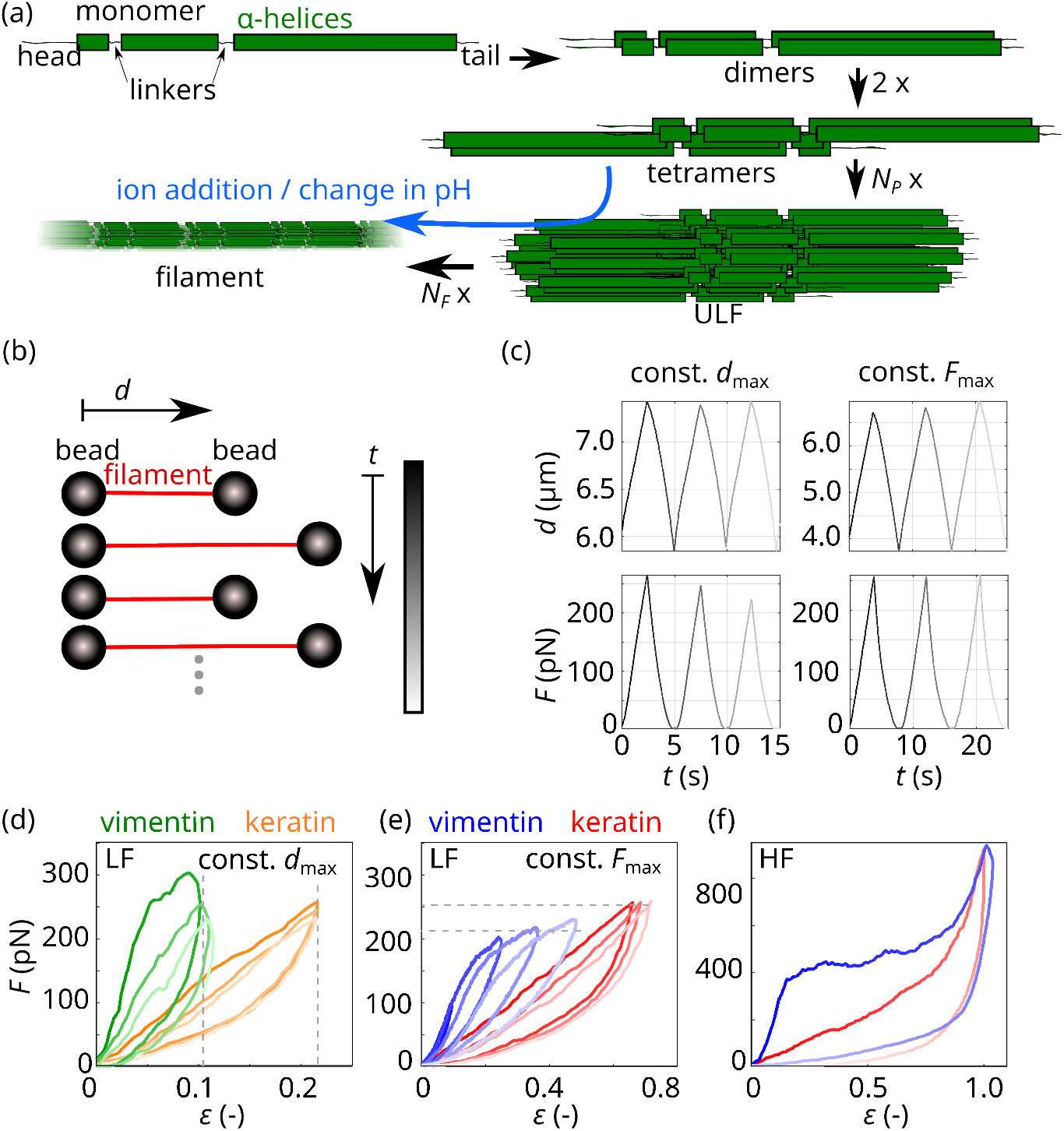
(a) Sketch of the assembly pathway of intermediate filaments. (b) Sketch of the measurement procedure. (c) Typical distance-time and force-time curves for constant *d*_max_ and decreasing *F*_max_ (left panels) and constant *F*_max_ measurements with an increasing *d*_max_ (right panels). Increasing time is indicated with lighter color. (d-f) Typical force-strain curves of keratin (orange, red) and vimentin (green, blue) filaments when (d) repeatedly stretched to a constant *d*_max_ (dashed gray lines) in the LF range, (e) repeatedly stretched to a constant *F*_max_ (dashed gray lines) in the LF range and (f) stretched once to the HF range and relaxed.

It remains an open question, whether the different force-strain behaviors of vimentin versus keratin filaments also lead to differing abilities to dissipate energy and absorb mechanical stress. In the case of vimentin filaments, the dissipative properties have already been well investigated: repeated stretching and filament relaxation experiment have shown that the *α* helices unfold during the first stretching and form random coils afterwards, which are then cycled in length.^19^ This non-equilibrium unfolding of *α* helices allows for energy dissipation of about 80% of the input energy.

Here we compare stretch-relaxation cycles for vimentin and keratin filaments. We find that the filament types behave fundamentally differently: Vimentin filaments soften, but maintain their original length, whereas keratin filaments elongate, but keep their original stiffness. Interestingly, both filament types are able to dissipate a large portion of the input energy, but via completely different physical mechanisms. We model the mechanical properties of keratin filaments numerically and draw an analogy to the mechanical properties of metals, which, by means of delocalized electrons, are highly deformable at constant stiffness, although the material elongates when stretched.^22^

## Results

### Keratin filaments elongate upon repeated loading

We previously showed that vimentin filaments, when repeatedly stretched and relaxed, do not plastically elongate but keep their original length.^18^ Here, to compare this property in detail for keratin and vimentin filaments, we employ a four channel-microfluidic chip as described in Ref. 16. We repeatedly stretch single filaments to a constant maximum distance *d*_max_ as sketched in Fig. 1b. As a consequence, the maximum force *F*_max_ decreases with each stretching cycle, i.e., over time *t* (Fig. 1c, left panels). In the plots, progressing time is indicated by lighter color. To be able to compare different filaments, independent of their individual length, we calculate the strain *ε* = Δ*L/L*_0_, i.e., we normalize the filament length gained by stretching Δ*L* = (*L − L*_0_) by the original filament length *L*_0_ at 5 pN.^16,17^ Vimentin filaments are stretched to a *d*_max_ that corresponds to the beginning of the plateau-like regime of the measured filament (green in Fig. 1d). Thus, with this measurement protocol, we probe filament mechanics just before a majority of the *α* helices within vimentin filaments start to open.^17,18^ Keratin filaments do not exhibit a plateau so that we fix *F*_max_ to 250 pN, which we define as the *low force regime* (LF).

To analyze whether the filaments are plastically deformed, i.e., elongated, during repeated stretching, we calculate the filament elongation by extrapolating linear fits to the elastic stretching regime of force-strain curves between *F* = 100 pN and *F* = 150 pN (gray shaded area), indicated by solid black lines in Fig. 2a. The extrapolation of the linear fits to the *x* axis (dashed lines in Fig. 2a) provides the effective length *ε*_*e*_ of the filament. This effective length is calculated in units of strain and indicated by the solid circles. We observe that keratin filaments elongate by *ε*_*e*_ = 0.1 *−* 0.2 after eight stretching cycles as shown in Fig. 2b (orange) whereas vimentin filaments barely elongate (green). ^19^ The elongation of the keratin filaments is in line with our previous hypothesis of sliding subunits within these filaments.^16^ Such slid subunits do not experience a restoring force. On the contrary, vimentin filaments elongate only very little as their subunits barely slide.

**Figure 2:**
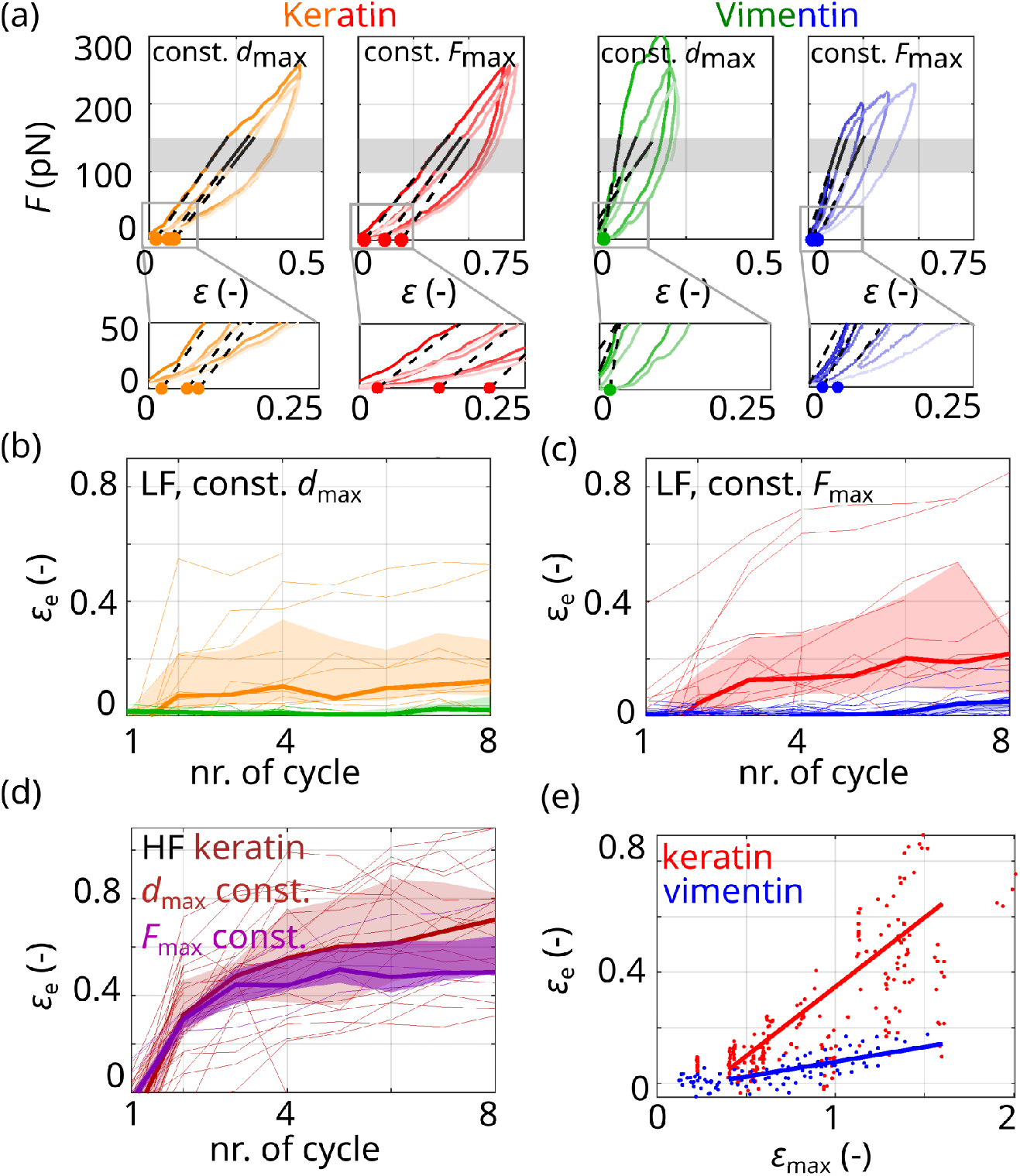
(a) Typical experimental data sets of keratin (orange, red) and vimentin (green, blue) filaments stretched three times to a constant *d*_max_ (orange, green) or to a constant *F*_max_ (red, blue) including fit curves to the force range of *F* = (100 − 150) pN (solid black lines). Fit ranges are indicated by gray shaded areas. Filament elongations (solid circles on the *x* axis) are determined by extrapolation of the linear fits to the *x* axis (dashed lines). (b,c) The effective length *ε*_*e*_ of filaments repeatedly stretched (b) to a constant *d*_max_ with *F*_max_ in the LF range or (c) to a constant *F*_max_ in the LF range. (d) *ε*_*e*_ of keratin filaments stretched to a constant *d*_max_ with *F*_max_ in the HF range (dark red) or to a constant *F*_max_ in the HF range (purple). (b-d) Thick lines show the median and shading indicates the area between the 25th and 75th percentile of the distributions per cycle. (e) *ε*_*e*_ of keratin (red) and vimentin filaments (blue) plotted against *ε*_max_ during the 4th to 15th cycle. The data are linearly fitted starting at a strain of 0.4 (solid lines).

Our first measurement protocol where we stretch the filaments to a fixed *d*_max_ for each cycle corresponds to biological situations where cells are elongated to a constant distance. However, there are also physiological settings, such as during muscle contraction, during which cells are repeatedly stretched to a constant force and we mimic this situation in our *in vitro* setting. Since keratin filaments do not exhibit a plateau-like regime, we chose a constant maximum force *F*_max_ = 250 pN that corresponds to the onset of the plateau-like regime in vimentin filaments, thus the onset of *α* helix unfolding. From a molecular point of view, this experimental protocol sheds light on the question of which mechanism within keratin filaments is instead responsible for elongation. We repeatedly stretch keratin and vimentin filaments to a constant maximum force *F*_max_ as shown in Fig. 1c, right panel. Consequently, the maximum distance increases from one cycle to the next. Typical force-strain curves are shown in Fig. 1e. We analyze the data in the same way as the data discussed so far, shown for keratin (red) and for vimentin (blue) in Fig. 2a. Compared to measurements where filaments are stretched to a constant maximum distance, we find that keratin filaments elongate further (compare red data in Fig. 2c to orange data in Fig. 2b). The increased elongation is likely caused by a further maximum distance during the stretching cycles due to a constant maximum force.

In specific biological situations, such as embryogenesis, cells may experience high forces and, consequently, high deformations. To include this *high force (HF) regime* in our study, and to test whether keratin filaments elongate further with higher loading forces, we stretch keratin and vimentin filaments to up to 900 pN and beyond the plateau-like regime of vimentin filaments as shown in Fig. 1f. Indeed, when stretched repeatedly to high forces, keratin filaments elongate by *ε*_*e*_ = 0.6 *−* 0.8 after eight cycles as shown in Fig. 2d for a constant maximum distance (dark red) and a constant maximum force (purple). To compare the different measurement protocols, we plot the effective length *ε*_*e*_ of the cycles 4 to 15 against the maximum applied strain *ε*_max_, see Fig. 2e. Under the high force regime, both types of filaments elongate, however keratin filaments (red) elongate further than vimentin filaments (blue). By fitting a linear relationship to the data above a strain of 0.4 (see red and blue solid lines in Fig. 2e), we estimate that keratin filaments elongate around five times more at a given maximum applied strain than vimentin filaments.

### Keratin filaments maintain their stiffness upon repeated loading

The pronounced elongation of keratin filaments during repeated loading raises the question of how this elongation impacts filament mechanics. For vimentin filaments, it is known that structural changes occur during repeated stretching and the filaments soften.^18,19^ We characterize the filament mechanics by determining their stretching stiffness *κ*_*f*_ from the linear fits shown in Fig. 2a. Remarkably, and in stark contrast to vimentin filaments,^18,19^ keratin filaments retain their stiffness, no matter whether they are pulled to a constant maximum distance or constant maximum force (orange and red in Fig. 3a and b, respectively). Likewise, when determining the filament stiffness from a fit to a fixed strain regime also results in a constant stiffness in keratin over all cycles (see discussion in the Supplementary Material). Thus, we can conclude that while keratin filaments are plastically deformed and their length “remembers” the loading history, their stiffness “forgets” it. Vimentin filaments behave in exactly the opposite way: They have a tensile memory concerning stiffness and “forget” the loading history with respect to their lengths. ^18,19^ Our results are confirmed when keratin filaments are stretched to the HF regime. Here as well, keratin filaments retain their stiffness while elongating during repeated pulling. Fig. 3c shows these data for a large maximum distance *d*_max_ (dark red) or high maximum force *F*_max_ (purple).

**Figure 3:**
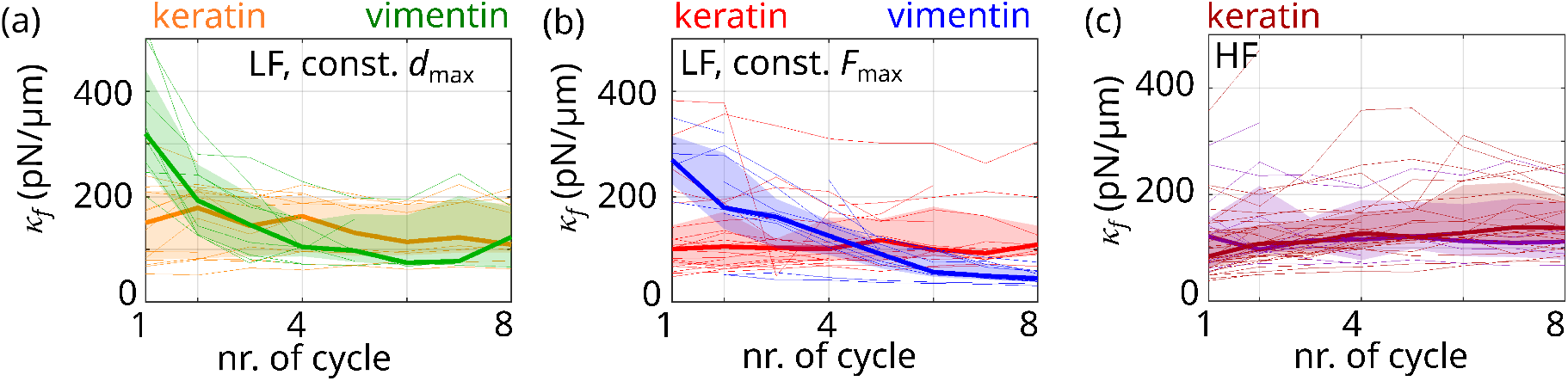
(a,b) Filament stiffness *κ*_*f*_ resulting from a fit to a force range of *F* = (100 − 150) pN (gray shaded area in Fig. 2a); (a) filaments stretched to a constant *d*_max_ (orange: keratin, green: vimentin); (b) filaments stretched to a constant *F*_max_ (red: keratin, blue: vimentin). (c) *κ*_*f*_ for keratin filaments stretched to a constant *d*_max_ in the HF range (dark red) and constant *F*_max_ in the HF range (purple). (a-c) Thick lines show the median and shading indicates the area between the 25th and 75th percentile of the distributions per cycle.

### An internal sliding mechanism explains the behavior of keratin filaments

The observation that the stiffness of keratin filaments is constant independent of the loading history and that the filament elongation depends on the loading history raises the question of which molecular mechanisms within keratin filaments cause this behavior that is so different from vimentin filaments. From repeated loading of vimentin filaments, we know that unfolded *α* helices do not directly transition to *β* sheets but turn into a third state, likely a random coil, which is softer than the *α* helices.^19^ Thus, if a significant portion of the *α* helices within keratin filaments was unfolded, we would expect a softening of repeatedly stretched filaments since the softer subunits within the filament would be stretched first. However, we observe a constant stiffness so that we conclude that most *α* helices within the keratin filament remain intact. We do not exclude the possibility that a small portion of *α* helices unfolds, but we hypothesize that these unfolded structures are not loaded during the next stretching cycle. Thus, we suggest that the following molecular mechanism accounts for our findings: The subunits within keratin filaments slide and form new bonds at a different location with the same properties as the original location as a consequence of the periodicity of the structure. Thus the stiffness during the next stretching cycle remains the same and bonds with the same properties as the previously existing bonds are stretched. The proposed sliding mechanism is sketched in Fig. 4a: Filament subunits are represented by gray rectangles. In case of keratin filaments, we assume that these subunits are dimers. These dimers are connected and interact as in a keratin filament in vitro (see Fig. 1a). Once a force is applied at time *t*_2_ *> t*_1_, the dimers can slide and new interactions can form (green arrows). One longitudinal and a lateral interaction has to break to allow for dimer sliding as shown in Fig. 4a, bottom. The sliding, longitudinally connected chain of dimers, i.e., the sliding “protofilament”, results in an elongation of the entire filament.^16^

**Figure 4:**
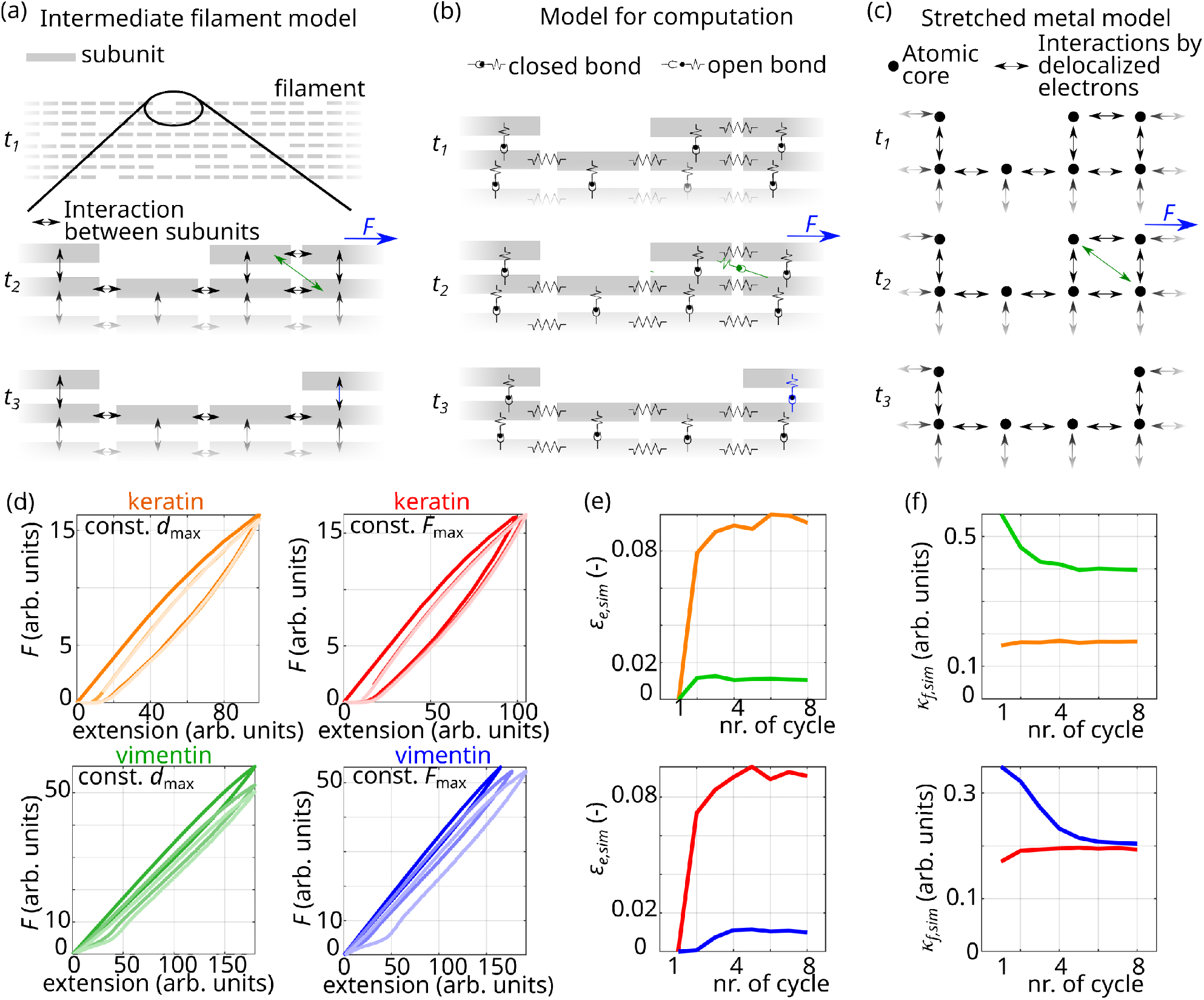
(a) Sketch of the proposed sliding mechanism in keratin filaments. The gray rectangles represent subunits such as dimers or tetramers. The arrows between the rectangles indicate interactions. The green arrows indicate the formation of new interactions. (b) Sketch of computational model to simulate force-extension curves of keratin filaments with the proposed sliding mechanism. The lateral interactions are modeled as elements which can either be in a closed or an open configuration. The longitudinal interactions are represented by springs. Under force at time *t*_2_ *> t*_1_, the lateral interactions break and subunits slide to the periodic position, where they form new lateral bonds (green). (c) Sketch of the sliding mechanism in metals in bulk. The black circles represent atoms, and the arrows show delocalized electrons which mediate the interactions between atoms. The shifted interactions are shown as green arrows. (d) Simulated force–strain curves of repeatedly stretched keratin filaments (top panel, orange/red) and vimentin filaments (bottom panel, green/blue); lighter colors represent progressing time. (e) Strain *E*_*e,sim*_ normalized by the maximum extension of stretched, simulated keratin filaments (orange, red) and stretched, simulated vimentin filaments (green, blue). (f) Stiffness *κ*_*f,sim*_ of stretched, simulated keratin filaments (orange, red) and stretched, simulated vimentin filaments (green, blue).

To show that this mechanism results in a constant stiffness, but further elongation of a filament upon repeated loading, we translate the suggested sliding mechanism into a computational model sketched in Fig. 4b. Interactions between dimers are represented by springs and elements which can unbind under force. A breaking interaction between two dimers is represented by an unbinding of these elements. Upon application of force, the dimers can slide and rebind (green elements in Fig. 4b, center). In our model, we assume that around 10% of dimers rebind to a neighboring dimer to approximate the experimental data. In this case, a defect arises at the position where the dimer was originally positioned (Fig. 4b, bottom). Slid dimers, which do not bind to a neighboring dimer, can rebind to the dimer which they were originally bound to. This sliding and rebinding mechanism is highly similar to the sliding of metallic atoms in a metal upon force application as sketched in Fig. 4c:^23^ Atoms are shown as black dots and interactions due to delocalized electrons are represented by arrows between the atoms. When a force is applied, the atoms slide and the electrons associate with a new atomic core (Fig. 4c, center). Just as our suggested model for keratin filaments, this sliding mechanism also results in an elongation of metals and a constant stiffness under repeated loading.^23^

Running the computational model for keratin filaments shown in Fig. 4b as a Monte-Carlo simulation, results in the force-extension curves shown in Fig. 4d, top panel. For comparison, a vimentin filament under repeated extension is modeled with the simulation presented in Ref. 18 (Fig. 4d, bottom panel). We analyze the stiffness of the filaments and the elongation in the same way we analyze the experimental data. In excellent agreement with the experiments, we find that keratin filaments retain their stiffness (orange, red in Fig. 4e) and vimentin filaments soften during repeated loading (green, blue). Keratin filaments extend (orange, red in Fig. 4f) while vimentin filaments return to their original length. Thus, instead of *α*-helical unfolding as in vimentin filaments, subunits slide within keratin filaments and thereby avoid major *α*-helical unfolding. Hence, next to crosslinkers,^19^ subunit sliding can protect *α* helices from unfolding.

### Keratin and vimentin filaments dissipate energy by different mechanisms

Vimentin filaments dissipate more than 80% of their input energy and there are strong indications that this occurs by by non-equilibrium *α*-helical unfolding.^18^ As keratin filaments do not posses this ability to unfold the *α* helices, the question remains if they dissipate a part of the input energy nevertheless, and if so, by which mechanism. To investigate this phenomenon in detail, we analyze the dissipated energy during their first stretching and relaxation cycle as shown in Fig. 5a and b as relative dissipated energy and absolute dissipated energy per filament length, respectively. We find that keratin filaments dissipate more than 50% of the input energy (red in Fig. 5a), which is lower than for vimentin filaments (blue), but still a considerable amount. Fig. 5b shows that in absolute units, both filament types dissipate energies on the order of 10^4^ *k*_*B*_*T/μ*m. The high amount of dissipated energy supports the notion that also keratin filaments may act as cellular shock absorbers.^18,24,25^

**Figure 5:**
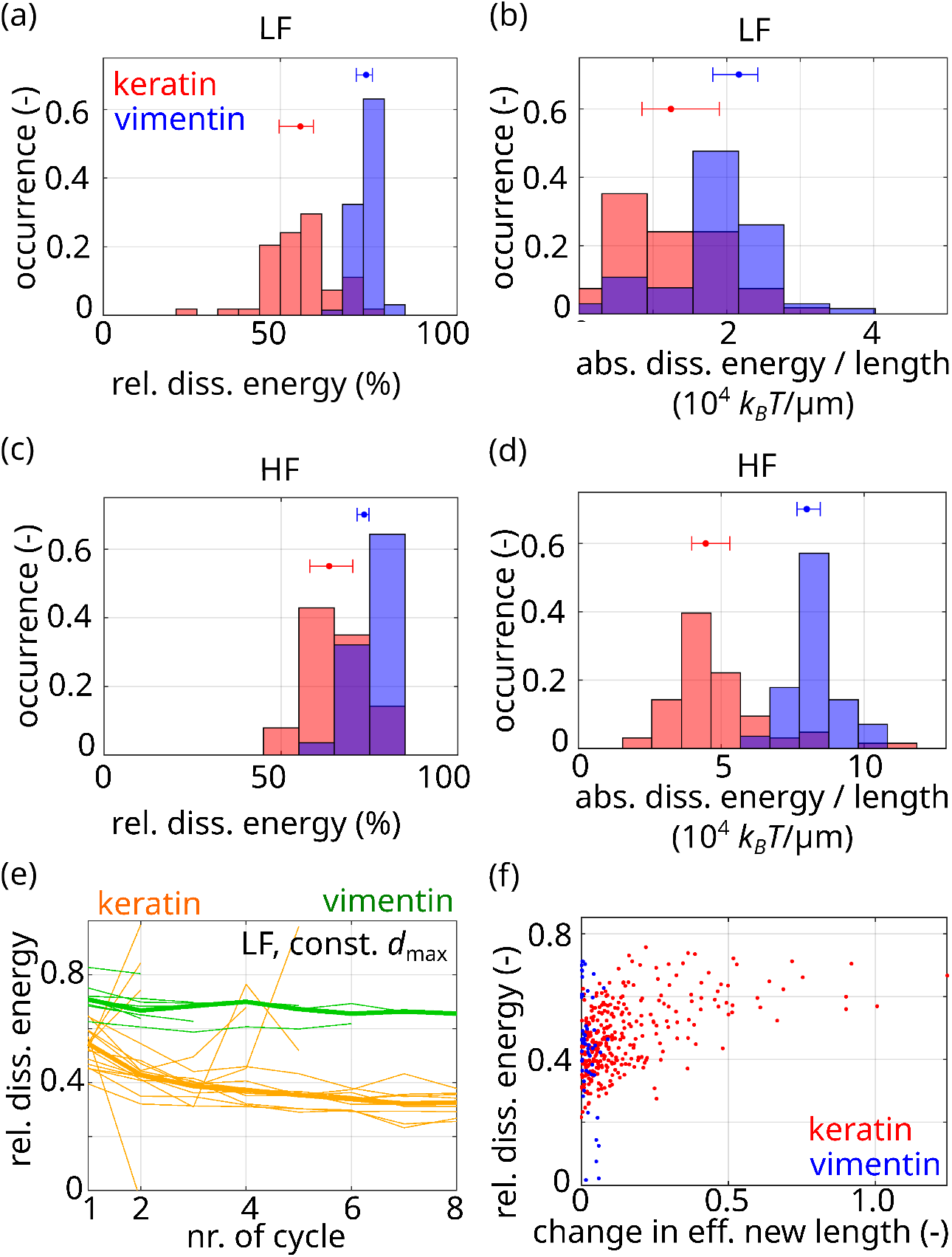
Dissipated energies of keratin (red) and vimentin filaments (blue) for the first stretching cycle. (a) Relative dissipated energy and (b) absolute dissipated energy per length of filaments stretched to the LF range. (c) Relative dissipated energy and (d) absolute dissipated energy per filament length when stretched to the HF range. Dots and whiskers indicate the median and the 25th and 75th percentile of the distributions, respectively. Bin width of the histograms is determined with the Freedman-Diaconis rule. (e) Relative dissipated energy plotted against the number of stretching cycles for keratin (orange) and vimentin (green) filaments stretched to a constant *d*_max_. (f) Relative dissipated energy of all cycles compared to Δ*ε*_*e*_.

Loading to higher forces (HF regime) also leads to high levels of relative energy dissipation (Fig. 5c) with around 60-70% of dissipated energy for keratin filaments (red) and 70-80% of dissipated energy of vimentin filaments (blue). The absolute dissipated energy increases when stretched to higher forces as shown in Fig. 5d. These results apply to the first stretching cycle. Considering further stretching of the filaments, we find that vimentin filaments dissipate about twice as much energy as keratin filaments when repeatedly stretched, see Fig. 5e.

Nevertheless, the question remains how energy is dissipated in keratin filaments on the molecular scale if it is not via the unfolding *α* helices. To dissipate energy, bonds need to be broken and must not rebind at the same position immediately. In the case of keratin filaments we show by our experiments and simulations that most bonds within the *α* helices remain intact, but the bonds between subunits are broken. Thus, we analyze the additional elongation Δ*ε*_*e*_ from one cycle to the next compared to the dissipated energy during that cycle, see Fig. 5f. The additional elongation is a measure for the number of slid dimers within the filament since dimer sliding causes elongation for keratin filaments. For keratin filaments (red in Fig. 5f), higher relative dissipated energies are correlated with a more pronounced increase of Δ*ε*_*e*_. Since vimentin filaments barely elongate, we do not observe such a correlation between the dissipated energy and Δ*ε*_*e*_ (blue in Fig. 5f). We therefore conclude that keratin filaments dissipate their energy by breaking bonds between subunits which results in subunit sliding and filament elongation, whereas vimentin filaments dissipate their energy by *α* helix unfolding.^18^

## Discussion and conclusion

Vimentin and keratin are an interesting pair of intermediate filaments as the “switch” between them plays a major role in the epithelial-to-mesenchymal transition, and therefore in cancer mestastasis, wound healing and embryogenesis. Here, by repeatedly loading of single keratin filaments and comparing the results to vimentin cycling, we find surprisingly different behaviors: Within keratin filaments, subunit sliding causes filament elongation and diminishes *α*-helical unfolding. As the *α* helices stay intact, the stiffness of keratin filaments remains constant independent of the loading history. Another consequence of the subunit sliding is keratin filament elongation. Contrarily, vimentin filaments retain their length but soften during repeated loading due to *α*-helical unfolding.^18,19^

Remarkably, both intermediate filament protein types possess the same secondary structure. The mechanical differences are routed in variations in the primary protein structure, i.e., the amino acid sequence. Differing charge and hydrophobicity patterns lead to different interactions within the filaments and cause the very distinct mechanical behaviors. ^16^ Specifically, a very distinct charge pattern of the vimentin monomer causes a tight arrangement, called compaction, of the vimentin tetramers within the filament.^26,27^ This compaction of vimentin filaments and stronger electrostatic and hydrophobic interactions within the filaments inhibit subunit sliding so that the *α* helix unfolding is energetically favorable with respect to subunit sliding. Keratin filaments do not exhibit this specific charge pattern necessary for compaction so that subunits can slide.^3,16,28,29^ For keratin, in contrast, we find that an analogy to the sliding mechanism of atoms within a metal: due to “sliding” of electrons within the periodic lattice of atom cores, metals change their length, but retain their stiffness and dissipate energy.^23^ In case of keratin intermediate filaments, subunits slide along one another other and can rebind due to the periodic structure of the filament, which results in a constant stiffness, elongation as well as energy dissipation. Thus, one may claim that keratin filaments mimic a biological metal-like response to mechanical stress. In contrast to keratin, we speculate that the stretching mechanism occuring in vimentin filaments might be analogous to the stretching of a double-network hydrogel:^30^ The tightly connected protofilaments correspond to the covalently crosslinked polymers ensuring stability of the gel, and the repeated breakage of bonds within the *α* helix or random coil correspond to the breakage of ionic crosslinks within the hydrogel, which dissipate energy.

Thus, the differential expression of intermediate filament proteins might be a way for cells to ensure that filaments either keep their original stiffness (keratin) or their original length (vimentin). We speculate that a constant stiffness during repeated loading can be a vital property for specific cell types such as skin cells so that they keep a constant pressure against the same applied force, but are flexible in their length. A constant filament length might be desirable for more motile cells so that cells or part of the cell are protected from elongation. A prominent example is the vimentin “cage” found surrounding the nucleus of cells.^31–33^ This is in line with the expression of vimentin in motile cells.^34^ Additionally, we find that vimentin dissipates considerably more energy even after repeated loading. Thus, motile cells might rather express vimentin instead of keratin to dissipate more energy caused by deformation due to the cellular movement. Similarly, the properties of vimentin filaments and their networks might be important during intracellular transport of large organelles: vimentin filaments form a dense network with a small mesh size, which needs to be deformable without permanently changing its structure, when cargo is transported through.

To conclude, our findings foster the idea of differential expression of intermediate filament proteins as a tool for cells to adapt their mechanical properties to their surrounding environment: After repeated loading, keratin filaments elongate, but exhibit a constant stiffness, while vimentin filaments retain their length and soften – although both filament types consist of monomers with the same secondary structure. We propose that weaker interaction strengths within keratin filaments than within vimentin filaments cause these distinct behaviors, because they allow for subunit sliding and thereby and protect the *α* helices within keratin intermediate filaments from unfolding. Interestingly, independent of the interaction strength within the two different filament types, both may act as cellular shock absorbers as they dissipate a major part of the input energy. Yet, the mechanism by which energy is dissipated is completely different and relies on internal viscous friction for keratin filaments and non-equilibrium unfolding of *α* helices for vimentin filaments.

## Acknowledgements

We thank S. Bauch for the purification of the proteins. We are grateful for fruitful discussions with J. Kraxner, A. V. Schepers and D. A. Weitz. The work was financially supported by the European Research Council (ERC, Grant No. CoG 724932) and the Studienstiftung des deutschen Volkes e.V.

