## Supplementary material for "Keratin filament mechanics and energy dissipation are determined by metal-like plasticity": SI

### Materials and Methods

**Protein expression, labeling and dialysis:** Protein is expressed, labeled and dialyzed as described in Ref. 1. After assembly, keratin filaments are diluted 1:10 with keratin assembly buffer (10 mM TRIS, *p*H 7.5)[2–4] and stored at 37°C for up to two days. Vimentin filaments are stored in vimentin assembly buffer (100 mM KCl, 2 mM phosphate buffer, *p*H 7.5)[5,6] at 4°. For measurements with the optical tweezers setup (LUMICKS, Amsterdam, Netherlands),

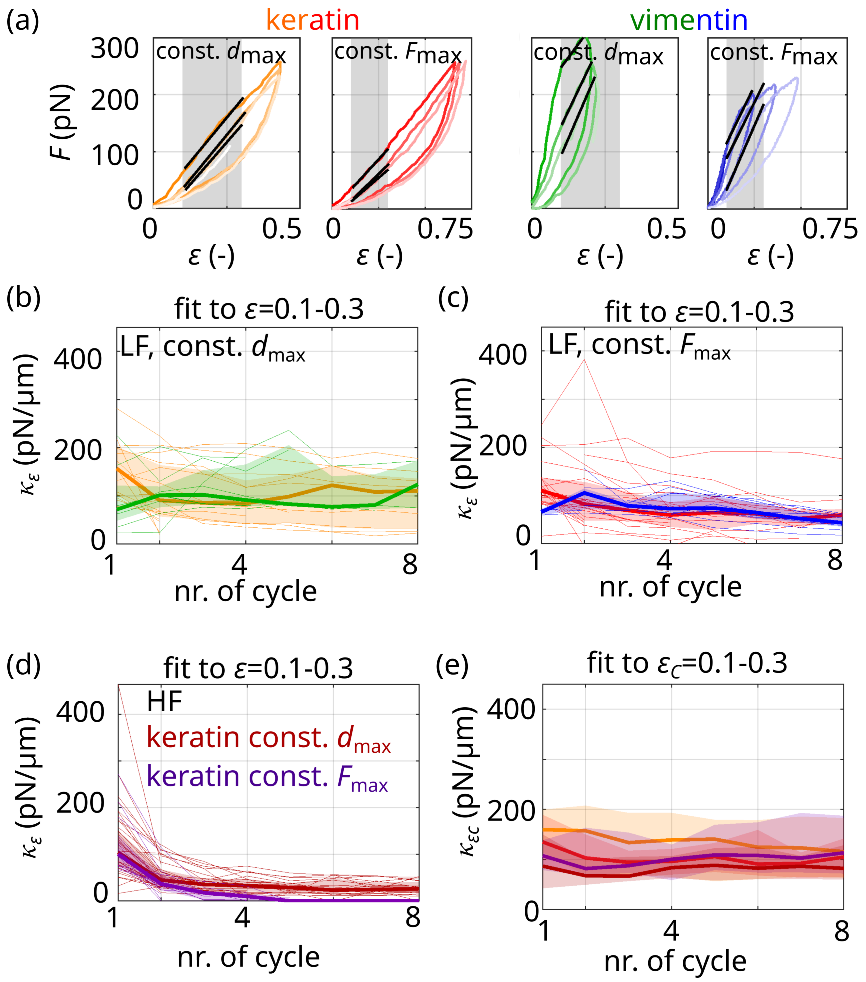

Figure S1: (a) Typical experimental data sets of keratin (orange, red) and vimentin (green, blue) filaments stretched three times to a constant $d_{\max}$ (orange, green) and to a constant $F_{\max}$ (red, blue) including fits to $\varepsilon = 0.1 - 0.3$ (gray shaded area, solid black lines); same data as shown in Fig. 2a in the main text. (b,c) Keratin (orange, red) and vimentin (green, blue) filament stretching stiffness $\kappa_\varepsilon$ resulting from the fits shown in a. (b) constant $d_{\max}$ with $F_{\max}$ in the LF range; (c) constant $F_{\max}$ in the LF range. (d) Keratin filament stiffness $\kappa_\epsilon$ derived from linear fits to $\varepsilon = 0.1 - 0.3$ for keratin filaments stretched to the HF range with a constant $d_{\max}$ (dark red) or to a constant $F_{\max}$ (purple). (e) Corrected $\kappa_{\epsilon c}$ for keratin filaments for all conditions studied: constant $d_{\max}$ with $F_{\max}$ in the LF range (orange), constant $F_{\max}$ in the LF range (red), constant $d_{\max}$ with a $F_{\max}$ in the HF range (dark red) and constant $F_{\max}$ in the HF range (purple). (b-e) Thick lines show the median and shading indicates the area between the 25th and 75th percentile of the distributions per cycle.

we load a four-channel microfluidic chip. Two maleimide-coated polystyrene beads (Kisker Biotech, Steinfurt, Germany) are captured with the optical tweezers from the beads channel and calibrated via the thermal noise spectrum. All filaments are stretched in vimentin assembly buffer, which corresponds to the high-ionic strength buffer condition used in Ref. 1.

All data analysis and image processing is carried out with self-written MatLab codes.

### Stiffness determined from fit to strain range

In the main text, we show the determination of the filament stiffness $\kappa_f$ form a linear fit to the force range of 100-150 pN. The filament stiffness $\kappa_\epsilon$ can also be obtained from a linear fit to a defined strain range, e.g. 0.1-0.3 as shown in Fig. S1a. This fit reveals a slight decrease of filament stiffness for both keratin and vimentin filaments (see Fig. S1b-d). However, since keratin filaments elongate, the fit range of the strain needs to be adjusted to the effective length $\varepsilon_e$ of the filament. We therefore calculate a corrected strain $\varepsilon_c = \varepsilon - \varepsilon_e$ for each cycle. With this corrected strain, we confirm the result above that the stiffness of keratin filaments is constant, see Fig. S1e.

### Simulations

Single keratin filaments under tension are modeled based on Refs. 1,7,8. One unit length filament (ULF) has $N_P$ monomers, which are arranged in parallel and all have the same length before loading. The monomers can be divided into $N_C$ $N_M$-mers, i.e., a keratin filament has $N_C = 8$ dimers with $N_M = 2$.[9] We refer to these $N_M$-mers as "subunits", which have the ability to slide with respect to each other. When subunits slide, the bonds between them open and re-form at the neighboring periodic position in the direction of the applied force as sketched in Fig. 4c in the main text. We assume that interactions within subunits are strong compared to interactions between these subunits so that the interactions within subunits, i.e. between monomers, do not break. Yet, the model is still valid if subunits are assumed to be equivalent monomers. Once a dimer has slid, it either pulls all dimers in a protofilament with it or it leaves a defect, i.e., an empty dimer as sketched in Fig. 4a,b in the main part.

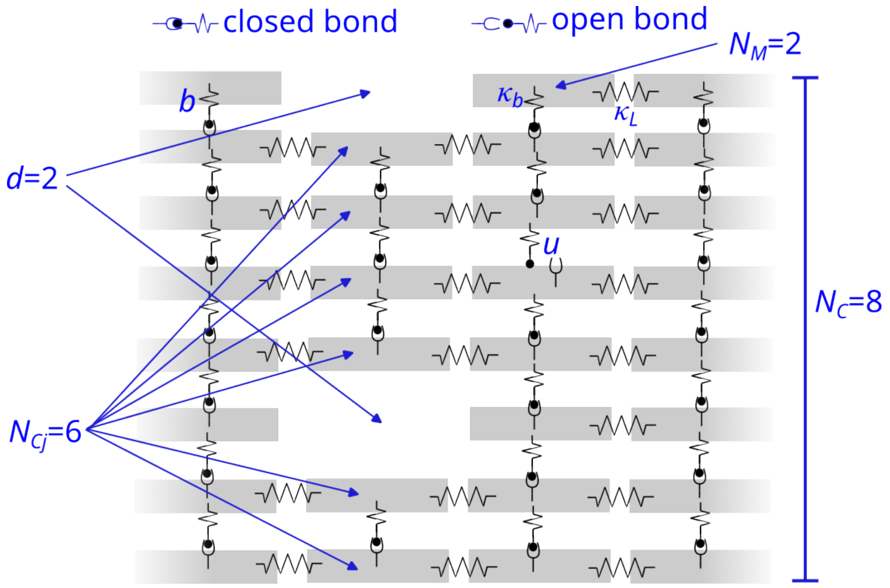


Figure S2: Sketch of the model describing subunit sliding.

We model the interaction between subunits by a spring with a spring constant $\kappa_b$ and by an element that can open into an unbound state $u$ when force is applied and can close again into a bound state $b$, when the connection to a new dimer forms. A sketch of the model is shown in Fig. S2. $B_j$ is the number of intact lateral bonds between dimers in the $j$th ULF. The longitudinal interaction between subunits forming a protofilament is described with a spring constant $\kappa_L$.

Thus, the spring constant of the $j$th ULF is:

$$\kappa_j = N_C \left( \frac{1}{\kappa_L} + \frac{1}{B_j \kappa_b} \right)^{-1} .$$

The spring constant of the entire filament is calculated by $\kappa_F = 1/\left(\sum_{j=1}^{N_F} 1/\kappa_j\right)$, where $N_F$ is the number of ULFs in a filament. Similar to Ref. 7, we neglect viscous and entropic contributions for simplicity.

We assume that the sliding of the subunits is a close-to-equilibrium process under force.

The equilibrium reaction constant $K_{eq}$ for subunits to unbind is:

$$K_{eq} = \frac{r^{b\to u}}{r^{u\to b}} = \exp(-\Delta G/(k_B T)) = 1/\gamma\,, \tag{1}$$

where we define $\gamma = \exp(\Delta G/(k_B T))$.

The force is distributed among the number of independent subunits $N_C$. The effect of the force on the rates is distributed between the binding and unbinding rates with a load distribution factor $\theta$,[10,11] which ensures detailed balance and thus that the model leads to a thermodynamic equilibrium state in the non-driven limit.

The force $\phi = F/F_b$ is dimensionless and scaled to the force $F_b$ which is required to open a bond between two subunits. The time $\tau$ is dimensionless and related to the time $t$ with the zero-force reaction rate from a subunit binding to another subunit to the unbound state $r_0^{b\to u}$ by $\tau = r_0^{b\to u} t$. We assume Bell-Evans kinetics,[12] so that the bound to unbound transition rate is:

$$r_{B_j}^{b\to u} = B_j r_0^{b\to u} \exp\left(\frac{\theta\phi}{B_j}\right). \tag{2}$$

In this study, we set $\theta = 1$, so only the unbinding rate is affected by the force. We assume that there is no restoring force acting on subunits which rebind to a neighboring subunit in the direction of loading as sketched in Fig. 4a,b in the main text. Yet, subunits can unbind without rebinding to the next subunit in the direction of loading because the disposition of the dimer is not sufficient to displace the subunit up to the periodic position. To describe the rebinding event to a new subunit or rebinding to the original subunit, we calculate the rebinding rate $r_{B_j}^{u\to b}$ depending on the force distributed on the $N_{Cj}$ subunits of the $j$th ULF (if the ULF possesses $d$ defects, the number of subunits $N_{Cj} = N_C - d$):

$$r_{B_j}^{u\to b} = r_0^{b\to u}\gamma \exp\left(\frac{-(1-\theta)\phi}{N_{Cj}}\right) \tag{3}$$

We assume that the filament posses 50 defects before it is stretched to approximate the experimental data. From these rates, we calculate the probability $P_{B_j}$ that a certain number of subunits is bound laterally to another subunit. The dynamics of this quantity are given by a Master equation 4 which we simulate using the Gillespie algorithm:

$$\begin{aligned} \frac{dP_{B_j}}{dt} &= r^{b\to u}_{B_j+1} P_{B_j+1} + r^{u\to b}_{B_j-1} P_{B_j-1} \\ &\quad -(r^{b\to u}_{B_{j,m}} + r^{u\to b}_{B_j}) P_{B_j}. \end{aligned} \tag{4}$$

When a subunit unbinds, the ULF extends by $\Delta L$ and with a probability of $p_S = 10\%$ the dimer slides and can bind to a new dimer to approximate the experimental data. If the filament is not stretched further than an effective elongation of 0.15, dimers do not slide and only unbind and rebind at the same position. We calculate the effective elongation by analyzing the extension of the filament during a retraction cycle at a force of 0.5 (arbitrary units).

To make the simulation run dimensionless, we normalize $\lambda = \Delta L / L_1$, where $L_1$ is a characteristic length of a unit-length filament. Thus, in this case, the extension of the $j$th ULF $\lambda_j$ is:

$$\lambda_j = \begin{cases} 0 & \text{if } B_j > 0 \\ 1 & \text{if } B_j = 0. \end{cases} \tag{5}$$

The total extension of the filament then is $\lambda_{tot} = \sum_{i=1}^{N_F} \lambda_j$. Since the optical traps pull on the filament with a constant velocity $v$, the end-to-end distance is $x(t) = vt$. The force on the filament becomes

$$\phi = \kappa_F (x - \lambda_{tot})\,. \tag{6}$$

To simulate the data shown in the main text in Fig. 6d-f, we use the following parameters:

Table 1: Dimensionless parameters and values for simulation of single stretched keratin filaments.

| Variable | Meaning | Value |
|---|---|---|
| $\Delta G$ | Energy difference between a laterally bound and unbound subunit | 6 |
| $\kappa_b$ | Stiffness of a lateral bond between two subunits | 4.6 |
| $\kappa_L$ | Stiffness of all linkers between subunits | 60 |
| $N_C$ | Number of subunits within a filament | 8 |
| $N_F$ | Number of unit-length filaments within a filament | 100 |
| $N_M$ | Number of monomers within a subunit | 2 |
| $N_P$ | Number of parallel monomers in a unit-length filament | 16 |
| $p_S$ | Probability that a slid dimer binds to the next dimer | 10% |
| $v$ | Loading speed | 10000 |

The simulation of single stretched vimentin filaments is from Ref. 7 with the following parameters:

Table 2: Dimensionless parameters and values for simulation of single stretched vimentin filaments.

| Variable | Meaning | Value |
|---|---|---|
| $\Delta G$ | Energy difference between a laterally bound and unbound subunit | 2 |
| $\kappa_\alpha$ | Stiffness of a lateral bond between two subunits | 5.5 |
| $\kappa_\beta$ | Stiffness of all linkers between subunits | 5.5 |
| $\kappa_L$ | Stiffness of all linkers between subunits | 100 |
| $N_F$ | Number of unit-length filaments within a filament | 100 |
| $N_P$ | Number of parallel monomers in a unit-length filament | 32 |
| $v$ | Loading speed | 2000 |

To convert these parameters into real values with actual units, fitting of the resulting force-strain curves would be necessary as shown in the Supplemental Material of Ref. 1.
